# Evaluating the potential role and contribution of transposable elements to the evolution of microbial multicellularity across the tree of eukaryotes

**DOI:** 10.64898/2026.06.24.734286

**Authors:** Angela Correa, Matthew W. Brown, Idan Banson, Robert E. Jones, Phigy Kalulu, Charles Thompson, Alexander K. Tice, David A. Ray

## Abstract

Multicellularity has evolved multiple times across the eukaryotic tree of life, including among protist lineages. Because transposable elements (TEs) strongly influence genome architecture and gene regulation, understanding their potential impact on genome structure and their relationship with gene expression may provide insight into the evolution of multicellularity.

Here, we generated a new genome assembly for the facultatively multicellular amoeba *Acrasis kona* and performed comparative analyses of TE composition, TE diversity, and TE-density organization across diverse protist lineages. Comparative analyses included unicellular and multicellular representatives from across the tree of eukaryotes, (Heterolobosea, Filasterea, Cristidiscoidea, and Chlorophyceae), including *Naegleria* spp., *Tetramitus jugosus*, *Capsaspora owczarzaki*, *Pigoraptor* spp., *Fonticula alba*, *Parvularia atlantis*, *Volvox carteri*, and *Chlamydomonas reinhardtii*.

To examine relationships between TEs and gene regulation, we integrated transcriptomic datasets from *A. kona*, *Capsaspora owczarzaki*, and *Volvox carteri* with genome-wide TE-density analyses of differentially expressed genes. TE abundance and composition varied substantially among lineages, with species that exhibit more complex developmental or cellular organization generally containing higher TE proportions than closely related unicellular taxa. Patterns of TE-density organization near up-regulated, down-regulated, and non-differentially expressed genes also differed among systems, ranging from strong TE depletion in *A. kona* to weaker or cell-type-specific patterns in *Capsaspora* and *Volvox*.

Together, these findings suggest that transposable elements are associated with multicellularity across diverse protist lineages, although the specific roles they play appear to be complex, lineage-specific, and not yet fully understood.

## Introduction

Transposable elements (TEs) are widely distributed genetic entities distinguished by their ability to move within and among genomes (Bourque et al., 2018). Through their capacity to relocate, TEs exert a substantial influence on genome composition, often accumulating over evolutionary time and playing a crucial role in shaping genome structure and function (Watson, 2014). Transposition itself is a powerful mutagenic process, but the presence of TEs can also modify genome structure by serving as substrates for non-homologous recombination and as sources of new genetic material (Wessler, 2006). Owing to their capacity to enhance genomic plasticity and variability, the study of TEs has become central to understanding how these elements contribute to species adaptation and evolutionary innovation (Casacuberta & González, 2013; Feschotte, 2008).

Because TEs can alter genome architecture, regulatory networks, and chromatin organization, their evolutionary impact becomes particularly relevant when investigating major transitions in biological complexity, including the evolution of multicellular lifestyles. The emergence of multicellular systems requires coordinated regulation of signaling, adhesion, communication, and differentiation pathways, processes that depend heavily on the organization and modulation of gene regulatory networks (King, 2004; Rokas, 2008). Increasing evidence indicates that TEs contribute to regulatory innovation by introducing novel cis-regulatory elements, modifying local chromatin environments, and influencing developmental gene expression patterns (Chuong et al., 2016; Sundaram & Wysocka, 2020). Consequently, studying TE landscapes may provide important insight into how genomes evolve to support increasingly complex developmental and cellular organizations.

Multicellularity is a complex trait that has independently emerged multiple times across the tree of life, including in fungi, diverse lineages of colonial and aggregatively multicellular protist lineages, charophyte green algae, the group from which land plants evolved, as well as other green, red, and brown algal lineages (Ruiz-Trillo et al., 2007). Over the past two decades, major advances in understanding the origins of multicellularity have been achieved through genomic comparisons among early-branching metazoans—such as cnidarians, ctenophores, and sponges (Putnam et al., 2007; Srivastava et al., 2008, 2010a; Moroz et al., 2014; Fortunato et al., 2014)—and their closest unicellular relatives within the Holozoa, including the choanoflagellates *Monosiga brevicollis* and *Salpingoeca rosetta* (King et al., 2008; Fairclough et al., 2013) and the filasterean *Capsaspora owczarzaki* (Suga et al., 2013; Sebé-Pedrós et al., 2017; Ocaña-Pallarès et al., 2022). Collectively, these studies have shown that multicellular traits frequently emerge through the modification and co-option of ancestral genes and regulatory pathways rather than through the origin of entirely novel gene repertoires, emphasizing the importance of comparative analyses among protists and other early-diverging eukaryotic lineages for understanding the evolution of biological complexity (Sebé-Pedrós et al., 2013, 2017; Ocaña-Pallarès et al., 2022).

Protists provide an especially valuable framework for investigating the evolution of multicellularity because they encompass a broad diversity of developmental and cellular organizations, ranging from strictly unicellular organisms to lineages capable of aggregation, cellular cooperation, or differentiated multicellularity (Sebé-Pedrós et al., 2017; Parfrey & Lahr, 2013). Comparisons among species sharing recent evolutionary histories but exhibiting contrasting developmental strategies therefore provide an opportunity to examine how changes in genome architecture, including TE composition and genomic distribution, may contribute to evolutionary diversification and increasing biological complexity. One important comparative system occurs within the volvocine green algae, which includes the unicellular species *Chlamydomonas reinhardtii* and the multicellular species *Volvox carteri*. Comparative genomic studies have demonstrated that *Volvox* retains most of the ancestral genes present in *Chlamydomonas*, while exhibiting regulatory modifications and expansions involving genes associated with extracellular matrix formation, developmental signaling, and cell-cycle regulation (Merchant et al., 2007; Prochnik et al., 2010; Herron, 2016). Transcriptomic analyses have additionally revealed extensive compartmentalization of gene expression between reproductive and somatic cell types in *Volvox carteri* (Klein et al., 2017), emphasizing the value of this system for investigating how TEs may relate to developmental gene regulation within differentiated multicellular organisms (Hanschen et al., 2017; Matt & Umen, 2018).

Comparable evolutionary systems are also found among heterolobosean and holozoan protists. The facultatively aggregative amoeba *Acrasis kona* alternates between unicellular growth and multicellular aggregation under specific environmental conditions, forming fruiting structures through coordinated morphogenesis (Brown, Silberman, & Spiegel, 2012). Recent genomic and transcriptomic analyses revealed that aggregation-associated genes in *A. kona* include homologs involved in signaling, adhesion, and differentiation pathways shared with other multicellular eukaryotes, suggesting that components of multicellular regulatory toolkits predate the emergence of complex multicellular organisms (Sheikh et al., 2024). Within Holozoa, the filasterean *Capsaspora owczarzaki* has become an important model for studying the origins of multicellular complexity because it exhibits distinct life stages and possesses genes associated with adhesion, signaling, and transcriptional regulation (Suga et al., 2013). More recently, comparative genomic analyses including *Pigoraptor chileana* and *Pigoraptor vietnamica* have highlighted substantial genomic and regulatory diversification among unicellular holozoans prior to the emergence of animals, emphasizing the importance of these lineages for understanding the evolutionary foundations of cellular and developmental complexity (Ocaña-Pallarès et al., 2022). Similar comparative frameworks are also represented within Cristidiscoidea by the aggregative multicellular species *Fonticula alba* and the unicellular relative *Parvularia atlantis*, which provide additional opportunities to investigate the evolutionary origins of multicellular organization across diverse protist lineages (Tice et al., 2022; López-Escardó et al., 2017).

Although previous studies have examined TE abundance and genome composition across diverse eukaryotic lineages, most comparative analyses have primarily focused on identifying TE classes and quantifying their genomic proportions using annotation approaches such as RepeatMasker. However, little is known about how TE density in protists differs across contrasting developmental strategies relative to up-regulated, down-regulated, and non-differentially expressed genes. This study addresses two overarching questions. First, do protist species exhibiting multicellular or aggregative lifestyles differ in TE abundance and diversity relative to unicellular representatives within the same major evolutionary lineages? Second, are up-regulated, down-regulated, and non-differentially expressed genes associated with distinct genomic TE-density environments across different protist systems? Based on current evidence regarding the influence of TEs on genome organization and regulatory evolution (Bourque et al., 2018; Chuong et al., 2016), we hypothesized that species exhibiting multicellular or aggregative lifestyles may exhibit increased TE abundance and distinct TE-density patterns surrounding up-regulated, down-regulated, and non-differentially expressed genes relative to related unicellular taxa.

To address these questions, we generated a new long-read genome assembly for *Acrasis kona* and performed comparative analyses of TE composition, diversity, and TE-density organization across representative species from Heterolobosea, Filasterea, Cristidiscoidea, and Chlorophyceae. Additionally, we integrated existing transcriptomic datasets from *A. kona*, *Capsaspora owczarzaki*, and *Volvox carteri* to evaluate whether developmentally regulated genes are associated with distinct genomic TE environments across multiple protist systems. These analyses provide a comparative framework for investigating how TE landscapes and TE-associated regulatory organization vary across lineages with distinct forms of developmental and cellular complexity.

## Methods

### Genome sequencing, assembly, and decontamination

High-molecular-weight genomic DNA from *Acrasis kona*, grown the yeast *Rhodotorula mucilaginosa* as a food source was sequenced using the Oxford Nanopore Technologies (ONT) PromethION P2 Solo platform. The data (∼14 Gb; read N50 ≈ 38 kb) was assembled de novo with Flye v2.9 (Kolmogorov et al., 2019), an assembler optimized for long-read ONT data, and subsequently polished with Medaka v1.7.3 to improve base-level accuracy. To detect putative contaminants, BLASTn v2.14 searches were performed against the previously published *A. kona* genome (Sheikh et al., 2024), the NCBI RefSeq bacterial database, and the *R. mucilaginosa* genome (GCA_034406715.1). Contigs matching bacterial or yeast sequences were removed using BEDTools v2.31.1, and sequences shorter than 10 kb were filtered with SeqKit v2.3.1.

Following decontamination, the filtered reads were reassembled with Flye to maximize contiguity and reduce fragmentation. Assembly quality was assessed with QUAST v5.2 (Gurevich et al., 2013), and completeness was evaluated using BUSCO v5.5.0 (Manni et al., 2021). The resulting polished and decontaminated assembly served as the reference genome for TE annotation, gene prediction, and TE-density analyses in this taxon. The *Acrasis kona* genome assembly has been deposited at NCBI under BioProject accession PRJNA1447523, BioSample accession SAMN57028165, and Whole Genome Shotgun accession JBWXZU000000000.

### Transposable element identification and TE library construction

Additional genome assemblies for comparative analyses were obtained from publicly available genomic repositories, including the NCBI Genome database and the European Nucleotide Archive (ENA) (Table 1). The dataset includes species exhibiting facultative or obligate multicellularity and their closely related unicellular relatives from the eukaryotic lineages Heterolobosea, Filasterea, Cristidiscoidea, and Chlorophyceae. This dataset allows for direct comparisons of TE proportion and diversity because these species share a relatively recent common ancestry but exhibit contrasting developmental or cellular organizations.

**Table 1.**
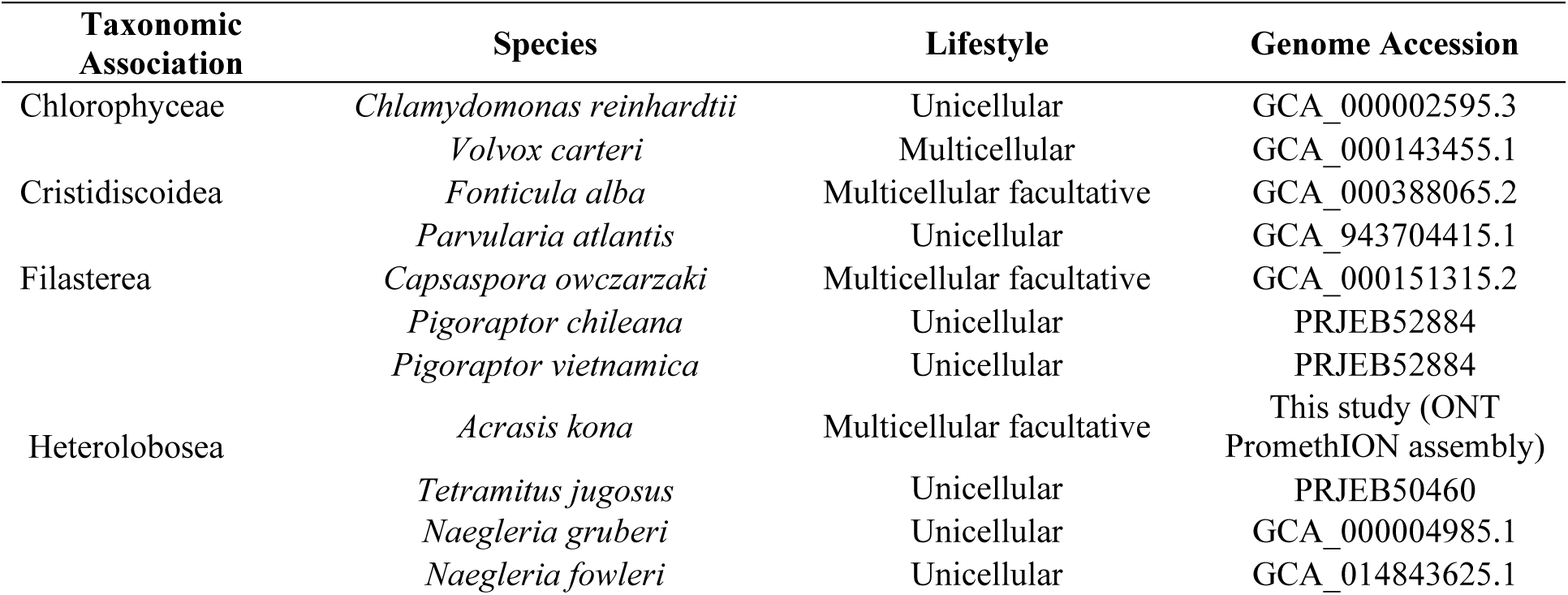
Genome assemblies analyzed in this study: Genome assemblies representing unicellular, multicellular, and facultatively multicellular protists were used to compare TE diversity and proportion across lineages. The *Acrasis kona* genome was newly assembled in this study from ONT PromethION data, while all other assemblies were retrieved from the NCBI Genome database. Accession numbers for each genome are provided.

To characterize transposable elements across genomes, a multi-tool detection strategy was used to increase sensitivity and classification accuracy. Five independent pipelines were executed: RepeatModeler2 (Flynn et al., 2020), MegaLTR (Mokhtar & El Allali, 2023), Inpactor2 (Orozco-Arias et al., 2018), HELIANO (Li et al., 2024), and HiTE (Hu et al., 2024). These tools combine structural and homology-based detection approaches and differ in their strengths, allowing comprehensive identification of both autonomous and non-autonomous elements. RepeatModeler2 provided broad de novo repeat discovery, while HiTE refined element boundaries. MegaLTR, Inpactor2, and HELIANO contributed additional class-specific resolution for LTR retrotransposons and Helitrons.

For each assembly the TE libraries generated by the five pipelines were merged and cross-compared. Redundant consensus sequences were removed by clustering with USEARCH (Edgar, 2010), resulting in a non-redundant, curated TE library. This library was used as the reference database for subsequent comparative analyses. RepeatMasker v4.1.5 (Smit, Hubley, and Green, 2013–2024) was then used to annotate each genome assembly using the curated TE library. Annotation outputs were parsed to quantify the proportion of masked genomic bases and to calculate the relative representation of major TE classes, including LTR retrotransposons, DNA transposons, Helitrons, LINEs, and SINEs.

### Gene prediction and functional annotation

Gene models for *Acrasis kona* were predicted using BRAKER3 (Gabriel et al., 2023) with soft-masked genome assemblies as input. Homology-guided predictions were generated using protein evidence from the UniProtKB Heterolobosea dataset, which integrates GeneMark-EP+ and AUGUSTUS. This gene set was used as the reference for all downstream analyses.

Functional annotation of *A. kona* genes was performed using a custom pipeline that combined sequence-similarity searches, domain prediction, and non-coding RNA detection. Predicted protein sequences from BRAKER3/AUGUSTUS were searched with BLASTp against a Heterolobosea UniProtKB protein database. BLASTp hits were filtered to retain matches with ≥60% identity, alignment lengths ≥50 amino acids, and e-values ≤1e-5. Conserved domains and Gene Ontology (GO) terms were further identified using InterProScan (Jones et al., 2014). Genes lacking significant BLASTp matches were additionally screened for non-coding RNA features using Infernal against the Rfam covariance model database using Rfam gathering thresholds for significant covariance-model matches (Nawrocki & Eddy, 2013). Annotation results were merged into downstream annotation tables and the BRAKER3 GFF3 file using custom Python and awk scripts rather than a packaged annotation merger. BLASTp- and InterProScan-derived annotations were linked back to BRAKER gene IDs after removing transcript suffixes, allowing transcript-level annotations to be reassigned at the gene level. For GFF3 integration, a custom Python script compared BRAKER3 gene coordinates with genes from the reference GFF3, normalized scaffold names, required same-strand overlap, and retained matches with at least 100 bp overlap and ≥50% reciprocal coverage between features. Functional fields, including gene name, locus tag, product description, and database cross-references, were then added as attributes to the corresponding BRAKER3 gene and transcript entries. This annotated gene set provided the basis for subsequent TE-density analyses.

### Transposable Element Density Analysis

The density of transposable elements in upstream regulatory regions was quantified using TE-Density v1.1.0 (Teresi, Teresi, & Edger, 2022). For each annotated gene, we calculated the proportion of bases annotated as TEs within fixed upstream windows (1 kb, 2 kb, and 5 kb) relative to the transcription start site. These distances were selected because they encompass the proximal promoter regions commonly associated with cis-regulatory activity in diverse eukaryotes.

TE-density analyses were conducted for *Acrasis kona*, *Capsaspora owczarzaki*, and *Volvox carteri*, the three species for which transcriptomic datasets representing distinct developmental or cellular conditions were available. TE-density estimates were subsequently integrated with differential expression analyses to evaluate whether developmentally regulated genes occur within distinct genomic TE environments. Genes were classified as upregulated, downregulated, or non-differentially expressed according to their corresponding transcriptomic comparisons, and TE-density distributions were compared among regulatory categories within each species.

### Differential Gene Expression Analysis

Differential gene expression analyses were conducted for *Acrasis kona*, *Capsaspora owczarzaki*, and *Volvox carteri* using publicly available RNA-seq datasets representing distinct developmental or cellular conditions. Raw sequencing reads were downloaded from the NCBI Sequence Read Archive (SRA) and reprocessed for alignment to their corresponding genome assemblies. Details regarding transcriptomic datasets and associated accession numbers are provided in Supplementary Table S1.

Read quality was assessed using FastQC v0.11.9 (Andrews, 2010), and adapter sequences and low-quality bases were removed using Trimmomatic v0.39 (Bolger, Lohse, & Usadel, 2014). Clean reads were aligned to their respective genomes using minimap2 v2.26 (Li, 2018) with splice-aware alignment settings to accommodate intron–exon structure. Gene-level read counts were obtained using featureCounts (Liao et al., 2014).

For *Acrasis kona*, differential gene expression during the transition between aggregation, growth, and germination conditions was analyzed using the RNA-seq datasets generated by Sheikh et al. (2024). Because these datasets lacked biological replicates, gene expression differences were estimated using GFOLD (Feng et al., 2012), consistent with the analytical approach implemented in Sheikh et al. (2024). Two pairwise comparisons were performed: aggregation versus growth and aggregation versus germination. Genes with positive GFOLD values were classified as upregulated, whereas genes with negative GFOLD values were classified as downregulated.

For *Capsaspora owczarzaki* and *Volvox carteri*, transcriptomic datasets with biological replicates were analyzed using DESeq2 v1.38.3 (Love, Huber, & Anders, 2014). In *C. owczarzaki*, differential expression analyses were performed across developmental life stages using RNA-seq datasets generated by Sebé-Pedrós et al. (2013). In *V. carteri*, transcriptomic comparisons between reproductive and somatic cell types were performed using RNA-seq datasets generated by Hanschen et al. (2017). Genes with adjusted p-values (padj < 0.05) were considered significantly differentially expressed and were subsequently classified as upregulated or downregulated according to log2 fold-change direction.

The resulting expression profiles for all three species were integrated with TE-density estimates to evaluate whether genes exhibiting distinct regulatory behaviors occur within different upstream TE-density environments relative to non-differentially expressed genes.

### Statistical Analysis of TE Density and Composition

Genome-wide TE proportion and composition were quantified using RepeatMasker annotations generated from the curated TE library produced by RepeatModeler2, MegaLTR, Inpactor2, HELIANO, and HiTE (Flynn et al., 2020; Orozco-Arias et al., 2018; Mokhtar & El Allali, 2023; Li et al., 2024; Hu et al., 2024). For each species, the total proportion of genomic bases masked as TEs and the relative contribution of major TE orders were calculated from RepeatMasker outputs, enabling comparative analyses across protist species exhibiting distinct developmental and cellular organizations.

To evaluate whether species exhibiting greater developmental or cellular complexity differ in TE proportion relative to unicellular representatives within the same evolutionary lineages, nonparametric analyses appropriate for small sample sizes and non-normal distributions were performed. Eight pairwise comparisons were conducted across four major protist lineages: Heterolobosea (*Acrasis kona* versus *Naegleria gruberi*, *N. fowleri*, *N. lovaniensis*, and *Tetramitus jugosus*), Filasterea (*Capsaspora owczarzaki* versus *Pigoraptor chileana* and *P. vietnamica*), Chlorophyceae (*Volvox carteri* versus *Chlamydomonas reinhardtii*), and Cristidiscoidea (*Fonticula alba* versus *Parvularia atlantis*). Paired differences in TE proportions were evaluated using a one-tailed Wilcoxon signed-rank test (Wilcoxon, 1945), testing the hypothesis that species exhibiting greater developmental complexity harbor higher overall TE proportions. Statistical robustness was additionally assessed using 10,000 paired sign-flip permutation replicates.

To quantify TE diversity within each genome, Shannon (H′) and Simpson (1–D) diversity indices were calculated from normalized TE-order proportions, summarizing both the richness and evenness of TE categories while accounting for differences in genome size. Principal component analysis (PCA) was performed using normalized TE-order proportions to evaluate similarities and differences in TE composition among protist genomes.

To evaluate relationships between TE density and gene regulation, TE-density estimates generated by TE-Density were integrated with differential expression analyses for *Acrasis kona*, *Capsaspora owczarzaki*, and *Volvox carteri*. Genes were classified as upregulated, downregulated, or non-differentially expressed according to their corresponding transcriptomic comparisons. For *A. kona*, analyses were conducted separately for aggregation versus growth and aggregation versus germination conditions. For *C. owczarzaki*, TE-density analyses were performed independently for aggregative versus adherent and filopodial versus aggregative developmental comparisons. For *V. carteri*, analyses compared reproductive and somatic cell-associated expression profiles.

Comparisons of upstream TE-density distributions (1 kb, 2 kb, and 5 kb windows) among regulatory categories were conducted using Mann–Whitney U tests and Welch’s t-tests. To further evaluate whether observed differences exceeded random expectations, permutation tests using 10,000 iterations were performed for all pairwise comparisons. Analyses were conducted both across all genes and across TE-positive genes only to evaluate whether the inclusion of genes lacking nearby TEs influenced observed regulatory patterns.

All statistical analyses were conducted in Python 3.10 using SciPy (Virtanen et al., 2020), Pandas (McKinney, 2010), NumPy (Harris et al., 2020), and scikit-learn (Pedregosa et al., 2011).

## Results

### Improved Genome Assembly of *Acrasis kona*

A new long-read genome assembly was generated for *Acrasis kona* using ONT PromethION sequencing, followed by a decontamination pipeline to remove bacterial, mitochondrial, and prey organism sequences identified during quality control. The resulting curated assembly shows substantial improvements in contiguity relative to the previously published version (Sheikh et al., 2024) (Table 2). Assembly contiguity increased markedly: N50 rose from 42.6 kb to 262.7 kb, and L50 decreased from 262 contigs to 47. The total number of contigs declined from 2,039 to 359, indicating a significantly less fragmented assembly. The total assembly size increased from 44.0 Mb to approximately 47.8 Mb, reflecting improved recovery of genomic regions while excluding contaminant bacterial contigs and separating mitochondrial and prey organism sequences. Ambiguous bases were nearly eliminated (7.56 to 0.21 per 100 kbp), further indicating improved assembly quality. BUSCO completeness remained stable after curation (85.5% vs. 85.1%), indicating that core gene content was preserved despite removal of contaminant sequences. However, the proportion of fragmented BUSCOs decreased (7.5% to 5.1%), suggesting improved gene contiguity, while duplicated BUSCOs increased slightly (10.6% to 12.2%), likely reflecting improved resolution of repetitive or paralogous regions. A small increase in missing BUSCOs (7.0% to 9.8%) is consistent with the removal of contaminant sequences that may have previously contributed spurious matches. Overall, these results indicate that assembly refinement substantially improved structural quality without compromising biological completeness.

**Table 2.**
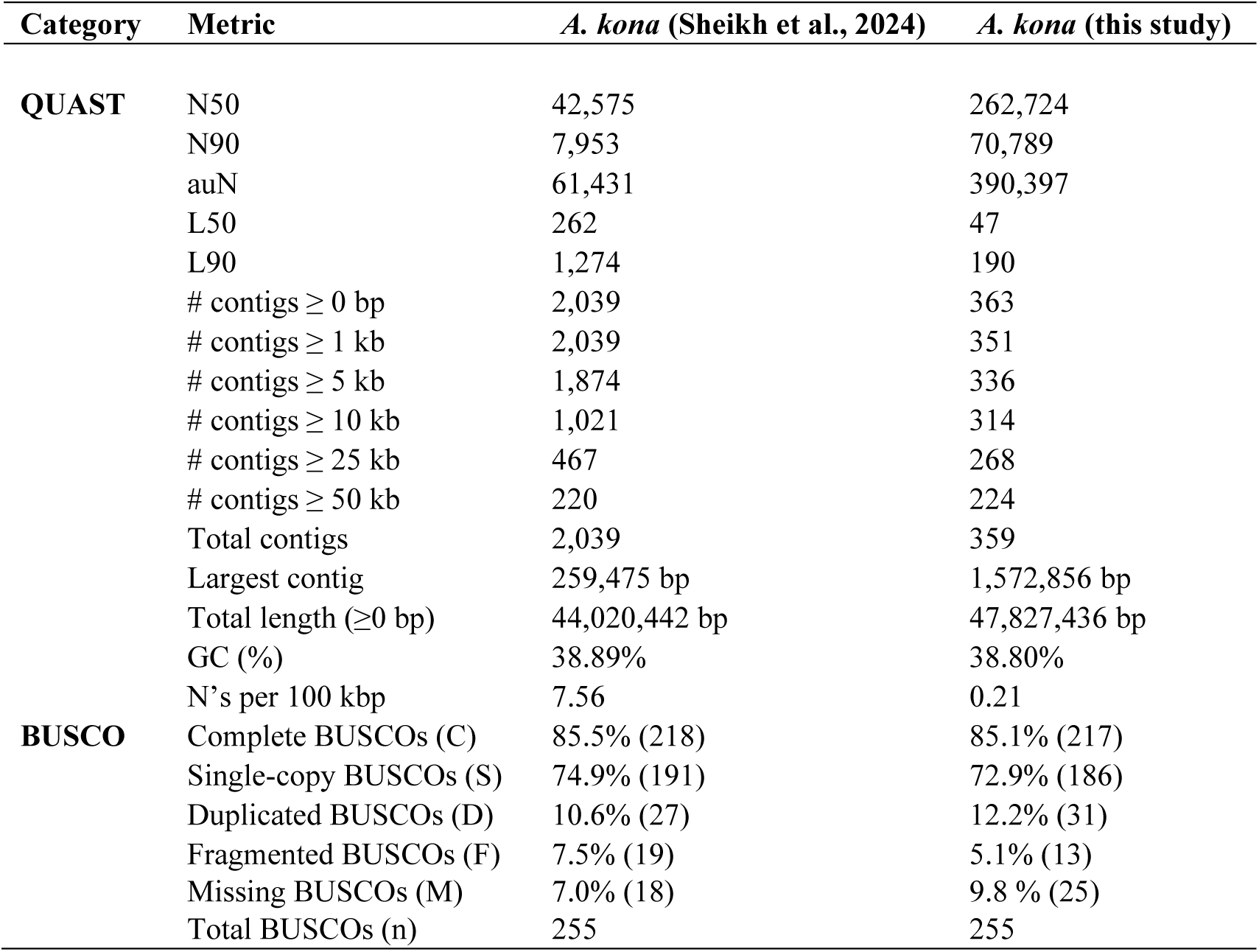
Assembly quality metrics for *Acrasis kona:* The original assembly from Sheikh et al. (2024) is compared with the new long-read assembly generated in this study. Metrics include contiguity statistics (N50, L50), total assembly length, GC content, ambiguous base frequency, and BUSCO completeness scores. The new assembly shows major improvements in contiguity and completeness, reflecting more accurate recovery of genomic structure.

Gene prediction and annotation for the curated *A. kona* assembly were performed using BRAKER3, which integrates RNA-seq evidence and protein homology to generate high-confidence gene models (Gabriel et al., 2023). The final annotation contained 25,067 predicted protein-coding genes after refinement and integration of transferred annotations from previous *A. kona* genome resources. Functional characterization was further improved using InterProScan, BLASTp searches against UniProtKB, and Infernal identification of non-coding RNAs, generating a consolidated annotation set for downstream comparative and TE-density analyses.

### Transposable Element Identification and Comparative Analysis

Using the curated, non-redundant TE library generated from RepeatModeler2, MegaLTR, Inpactor2, HELIANO, and HiTE, each protist genome assembly was screened with RepeatMasker to quantify the genomic proportion and class composition of transposable elements. Details regarding TE consensus sequences, classification, and identification methods are provided in Supplementary Table S2, and the complete curated TE library used for RepeatMasker analyses is provided as Supplementary File S1. This unified annotation strategy enabled consistent comparisons across representative protist species exhibiting contrasting developmental and cellular organizations.

Across the surveyed lineages, substantial variation in total TE proportion and TE composition was observed among species (Table 3; Figure 1). Among heteroloboseans, *Acrasis kona* contained 5.52% repetitive DNA, substantially exceeding the TE proportions detected in *Naegleria gruberi* (0.28%), *Naegleria fowleri* (0.28%), *Naegleria lovaniensis* (0.13%), and *Tetramitus jugosus* (0.14%). The TE landscape of *A. kona* was dominated by LTR retrotransposons (2.55%) and DNA transposons (2.00%), with additional contributions from rolling-circle Helitron elements (0.89%). In contrast, the compared heterolobosean unicellular species exhibited markedly reduced TE content across all TE categories. Within Filasterea, *Capsaspora owczarzaki* exhibited a total TE proportion of 3.86%, driven primarily by rolling-circle elements (1.94%) and DNA transposons (1.20%). Interestingly, the two *Pigoraptor* species displayed similarly elevated TE proportions, with *Pigoraptor chileana* and *Pigoraptor vietnamica* containing 3.18% and 3.24% repetitive content, respectively. In both *Pigoraptor* species, DNA transposons represented the dominant TE category, exceeding 2% of the genome. These results indicate that elevated TE abundance within Filasterea is not restricted to *Capsaspora* and may instead characterize multiple holozoan lineages exhibiting distinct developmental or cooperative cellular behaviors.

**Figure 1.**
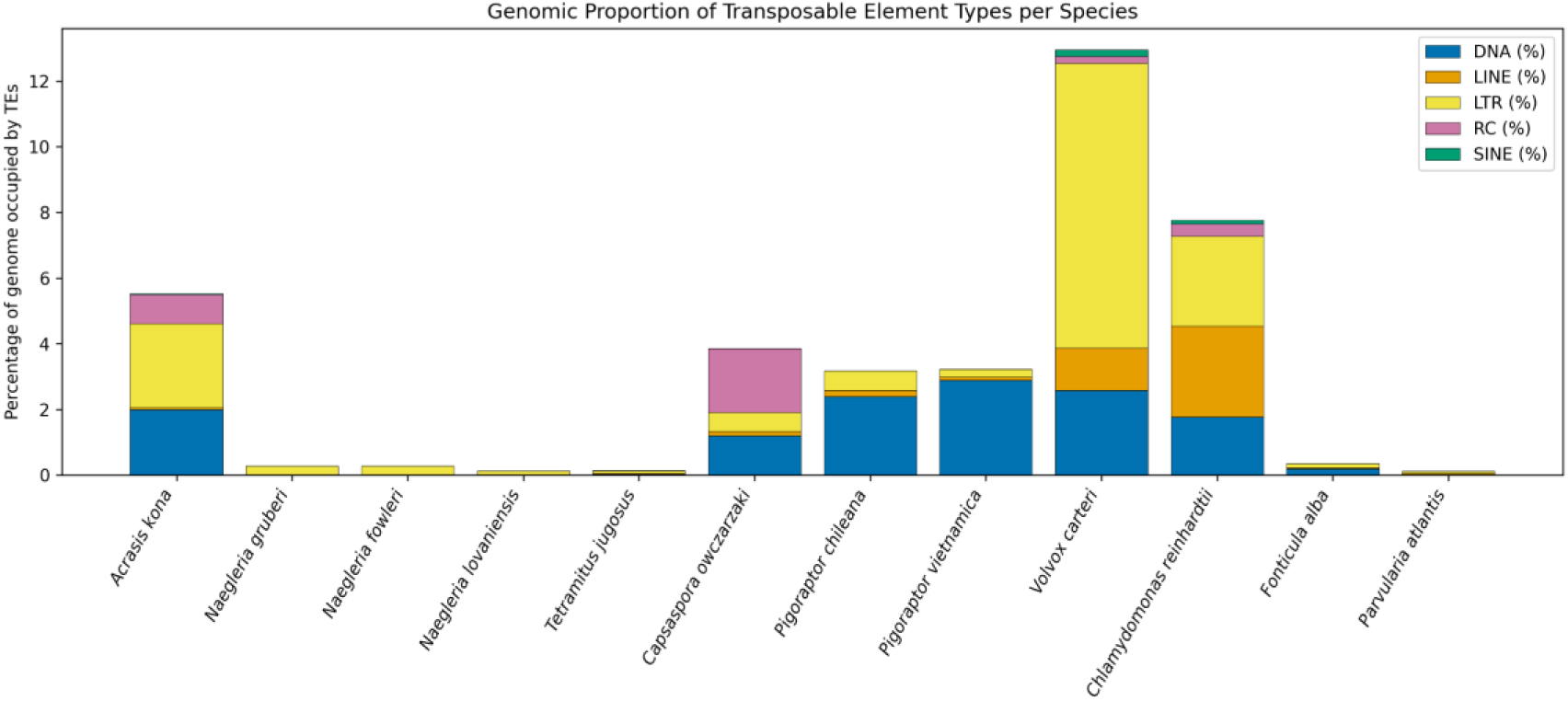
Genomic proportions of major transposable element types across paired protist species: The stacked barplot shows the percentage of each genome occupied by five major transposable element (TE) categories: DNA transposons, LINEs, LTR retrotransposons, rolling-circle (RC/Helitron) elements, and SINEs. Species are arranged along the x-axis according to major evolutionary lineages, including Heterolobosea, Filasterea, Chlorophyceae, and Cristidiscoidea. Considerable variation in both total TE proportion and TE composition was observed among species. In several lineages, including Heterolobosea and Chlorophyceae, aggregative or multicellular representatives exhibited substantially expanded TE landscapes relative to related unicellular species, particularly through the accumulation of LTR retrotransposons and DNA transposons. Within Filasterea, *Capsaspora owczarzaki* exhibited slightly higher total TE content than both *Pigoraptor* species, although all three genomes contained comparatively TE-rich landscapes dominated primarily by DNA transposons and rolling-circle elements.

**Table 3.**
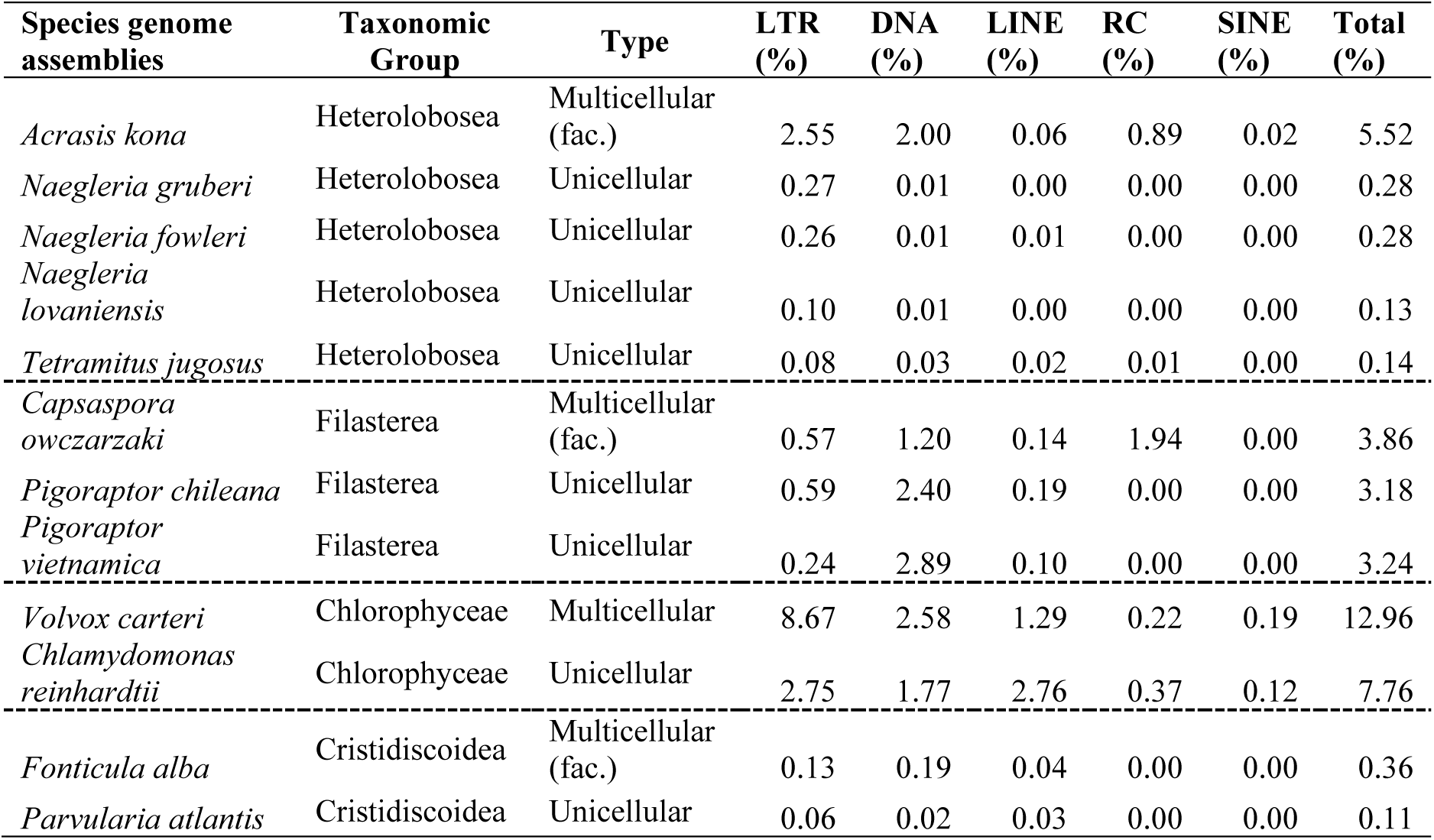
Proportion of transposable elements by class across unicellular and multicellular protist genomes: Proportional genomic content of major transposable element (TE) classes—including DNA transposons, LINEs, LTR retrotransposons, rolling-circle Helitrons, and SINEs—is shown for each unicellular and multicellular (or aggregatively multicellular) protist species analyzed. Values represent the percentage of the genome masked by each TE class based on RepeatMasker screening using the curated TE library generated in this study. The table highlights substantial lineage-specific variation in both total TE content and class composition across protist genomes.

Within Chlorophyceae, *Volvox carteri* exhibited the highest TE proportion among all analyzed species, with 12.96% of the genome annotated as repetitive DNA. This increase was largely driven by extensive accumulation of LTR retrotransposons (8.67%), accompanied by substantial DNA transposon (2.58%) and LINE (1.29%) content. In comparison, *Chlamydomonas reinhardtii* contained 7.76% repetitive DNA, with a more balanced TE composition distributed among LTR, LINE, and DNA transposon categories. Although both examined volvocine algae contained relatively TE-rich genomes, *V. carteri* exhibited substantially greater expansion of LTR retrotransposons. Within Cristidiscoidea, *Fonticula alba* contained 0.36% repetitive DNA, compared with 0.11% in *Parvularia atlantis*. Although both genomes exhibited comparatively low TE abundance, *F. alba* consistently displayed higher proportions across most TE categories.

Overall, these analyses revealed strong lineage-specific variation in TE composition and abundance across protist genomes. Multicellular or aggregative taxa generally tended to exhibit higher overall TE content, whereas unicellular representatives often displayed reduced and compositionally simpler TE landscapes. However, this pattern was not universal across all examined taxa, particularly within Filasterea, where elevated TE abundance was also observed in the *Pigoraptor* genomes, which are primarily unicellular but exhibit transient aggregative and multicellular-like behaviors under certain conditions (Hehenberger et al., 2017; Hehenberger et al., 2020).

To statistically evaluate whether species exhibiting greater developmental or cellular complexity consistently harbor increased TE proportions, paired nonparametric analyses were conducted across the eight phylogenetic comparisons representing Heterolobosea, Filasterea, Chlorophyceae, and Cristidiscoidea. The paired Wilcoxon signed-rank test revealed a significant enrichment of TE content among the species exhibiting greater developmental complexity (W = 36, one-tailed p = 0.0039). Similarly, paired sign-flip permutation analyses supported the robustness of this pattern (observed mean difference = 3.50; permutation p = 0.0053), indicating that the observed differences in TE proportion are unlikely to result from stochastic variation alone.

These results are summarized in Figure 1, which illustrates substantial differences in both total TE proportion and relative TE-class representation among the surveyed protist species. The stacked barplot highlights clear lineage-specific variation in the contribution of major TE categories, including LTR retrotransposons, DNA transposons, LINEs, rolling-circle elements, and SINEs.

TE diversity quantified using Shannon (H′) and Simpson (1–D) indices, revealed lineage-specific rather than universal patterns (Figure 2). Within Heterolobosea, *Acrasis kona* showed high TE diversity (H′ = 1.09; 1–D = 0.63), markedly exceeding the three *Naegleria* species, which exhibited low diversity values consistent with their reduced TE content. However, *Tetramitus jugosus* also showed relatively high diversity (H′ = 1.12; 1–D = 0.61), indicating that TE diversity within Heterolobosea does not strictly follow the same trend as total TE abundance. This pattern may partially reflect the influence of low overall TE content on diversity estimates, where relatively few TE families distributed across multiple categories can yield elevated diversity indices despite limited total TE abundance.

**Figure 2.**
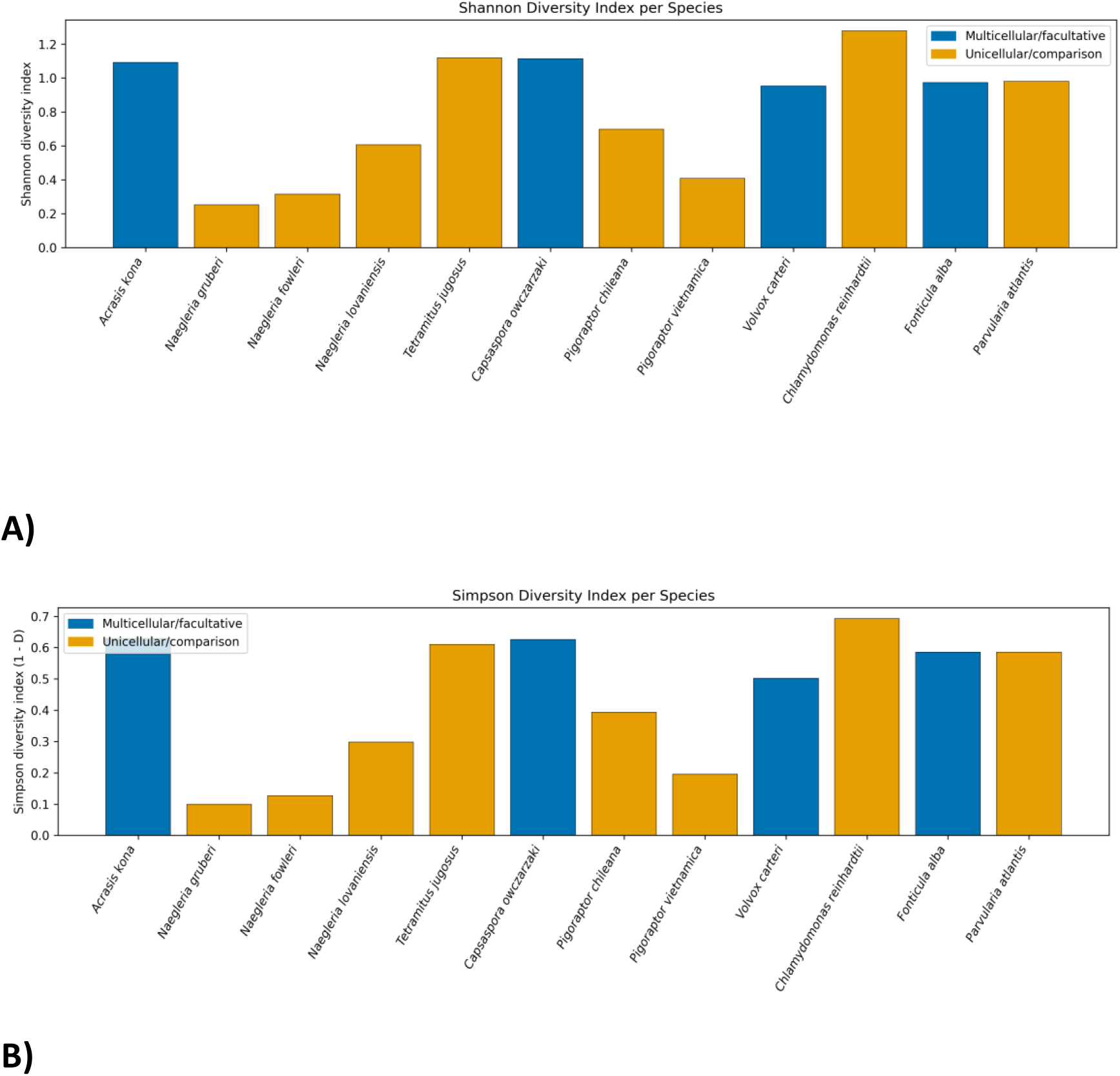
Transposable element diversity across multicellular and unicellular protist species: (A) Shannon diversity index (H′). (B) Simpson diversity index (1–D). Species are arranged according to major evolutionary lineages, with blue bars representing species exhibiting greater developmental or cellular complexity and orange bars representing unicellular comparison taxa. TE diversity exhibited strong lineage-specific variation rather than a universal relationship with multicellularity or aggregative behavior. *Acrasis kona* displayed substantially higher TE diversity than the compared *Naegleria* species, whereas *Tetramitus jugosus* also exhibited relatively high diversity despite its low overall TE proportion. Within Filasterea, *Capsaspora owczarzaki* exhibited greater TE diversity than both *Pigoraptor* species. In contrast, the unicellular alga *Chlamydomonas reinhardtii* exhibited the highest overall TE diversity among all analyzed genomes, despite containing lower total TE abundance than *Volvox carteri.* These results indicate that increased TE abundance does not necessarily correspond to increased TE diversity across protist lineages

Within Filasterea, *Capsaspora owczarzaki* exhibited higher TE diversity (H′ = 1.11; 1–D = 0.63) than both *Pigoraptor chileana* (H′ = 0.70; 1–D = 0.39) and *Pigoraptor vietnamica* (H′ = 0.41; 1–D = 0.20). In Chlorophyceae, *Chlamydomonas reinhardtii* had the highest overall TE diversity among all surveyed genomes (H′ = 1.28; 1–D = 0.69), despite having lower total TE abundance than *Volvox carteri*. In Cristidiscoidea, *Fonticula alba* and *Parvularia atlantis* showed nearly identical diversity values despite differing in total TE proportion.

A principal component analysis (PCA) based on proportional representation of TE classes (Figure 3) revealed substantial lineage-specific differences in TE composition among the surveyed protist genomes. PC1 explained 38.4% of the total variance, whereas PC2 accounted for an additional 30.9%, together capturing the majority of variation in TE composition across species. Separation along both axes reflected differences in the relative contribution of major TE classes, particularly LTR retrotransposons, DNA transposons, rolling-circle elements, and LINEs.

**Figure 3.**
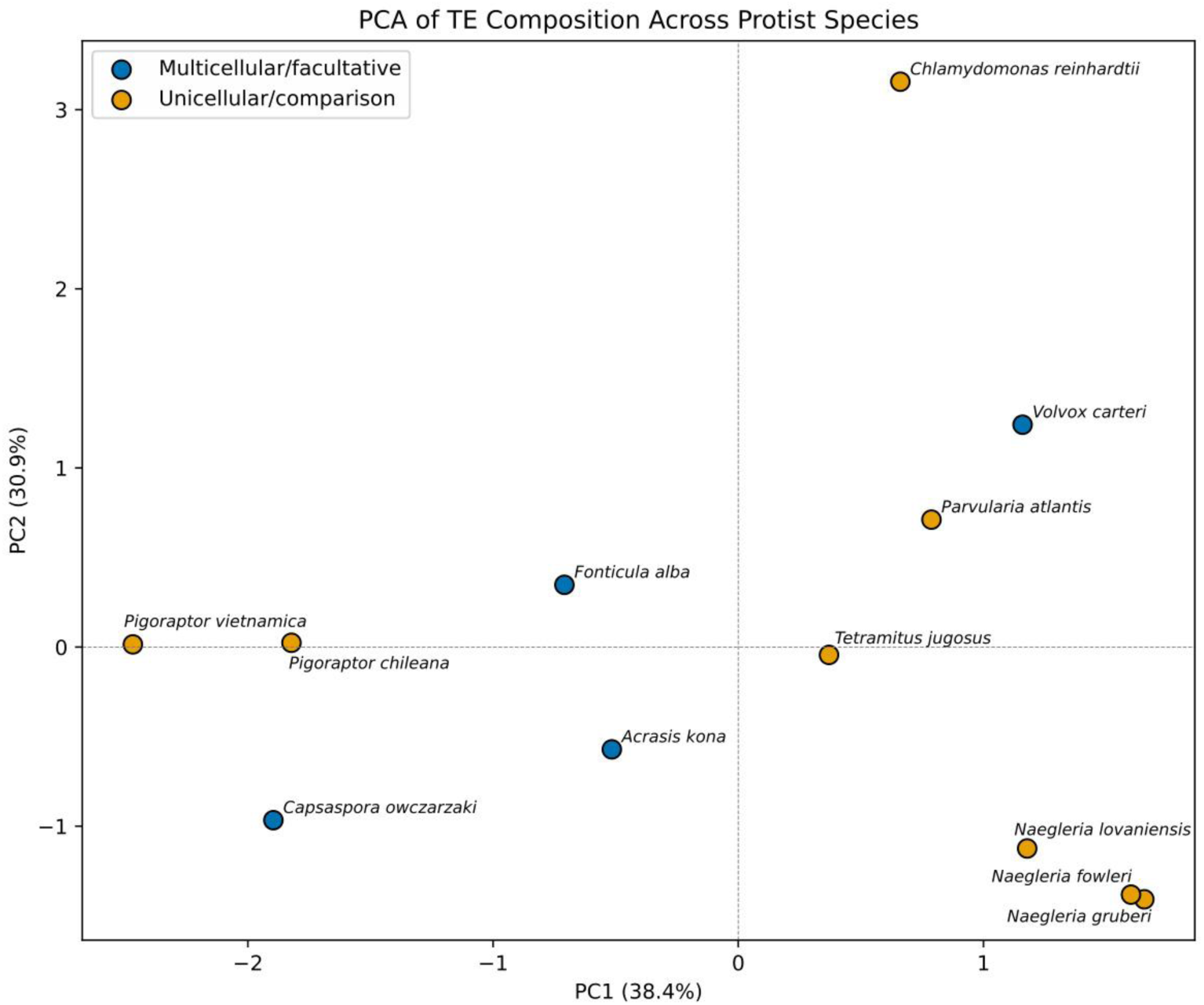
Principal component analysis (PCA) of transposable element composition across representative protist genomes. The PCA characterizes genome-wide TE composition using the proportional representation of major TE classes, including LTR retrotransposons, DNA transposons, LINEs, rolling-circle (RC/Helitron) elements, and SINEs. Blue points represent species exhibiting greater developmental or cellular complexity, whereas orange points represent unicellular comparison taxa. PC1 explains 38.4% of the total variance and PC2 explains 30.9%. Species are distributed according to lineage-specific differences in TE composition and relative TE-class abundance. *Acrasis kona* is separated from the *Naegleria* species primarily by its increased contribution of LTR retrotransposons and rolling-circle elements, whereas *Capsaspora owczarzaki* differs from the *Pigoraptor* species through greater RC/Helitron representation. Within Chlorophyceae, *Volvox carteri* and *Chlamydomonas reinhardtii* occupy distinct positions consistent with major differences in LTR retrotransposon abundance. Overall, the ordination highlights substantial variation in TE composition across protist lineages rather than a single conserved TE landscape associated with multicellularity or aggregative behavior.

Several multicellular or aggregative representatives occupied distinct regions of the ordination space relative to related unicellular taxa. *Acrasis kona* was clearly separated from the three *Naegleria* species along PC1, consistent with its expanded contribution of LTR retrotransposons and rolling-circle elements. Similarly, *Capsaspora owczarzaki* was differentiated from both *Pigoraptor* species, reflecting its greater relative abundance of rolling-circle elements. Within Chlorophyceae, *Volvox carteri* and *Chlamydomonas reinhardtii* were separated primarily along PC2, driven by the strong LTR retrotransposon enrichment observed in *V. carteri* and the comparatively balanced TE composition of *C. reinhardtii*.

Alternatively, several taxa occupied relatively similar regions of PCA space despite differences in total TE abundance. For example, *Fonticula alba* and *Parvularia atlantis* clustered relatively closely, suggesting similarities in proportional TE composition despite differing total TE proportions. In contrast, *Tetramitus jugosus* occupied a distinct position from the three *Naegleria* species despite belonging to the same major lineage and exhibiting comparatively low overall TE abundance. Although the PCA did not reveal a consistent clustering pattern associated with multicellular or aggregative lifestyles across the examined taxa, it did indicate that TE composition varies strongly among protist lineages and that increases in total TE abundance are often accompanied by shifts in the relative representation of specific TE categories rather than by uniform expansion across all TE classes.

### TE-density patterns across developmental regulatory groups

Comparisons of upstream TE density across developmental regulatory categories revealed substantial variation among the analyzed protist systems (Table 4). Patterns differed both in magnitude and direction, indicating that relationships between TE density and developmental gene regulation are highly lineage-specific rather than universally conserved.

**Table 4.**
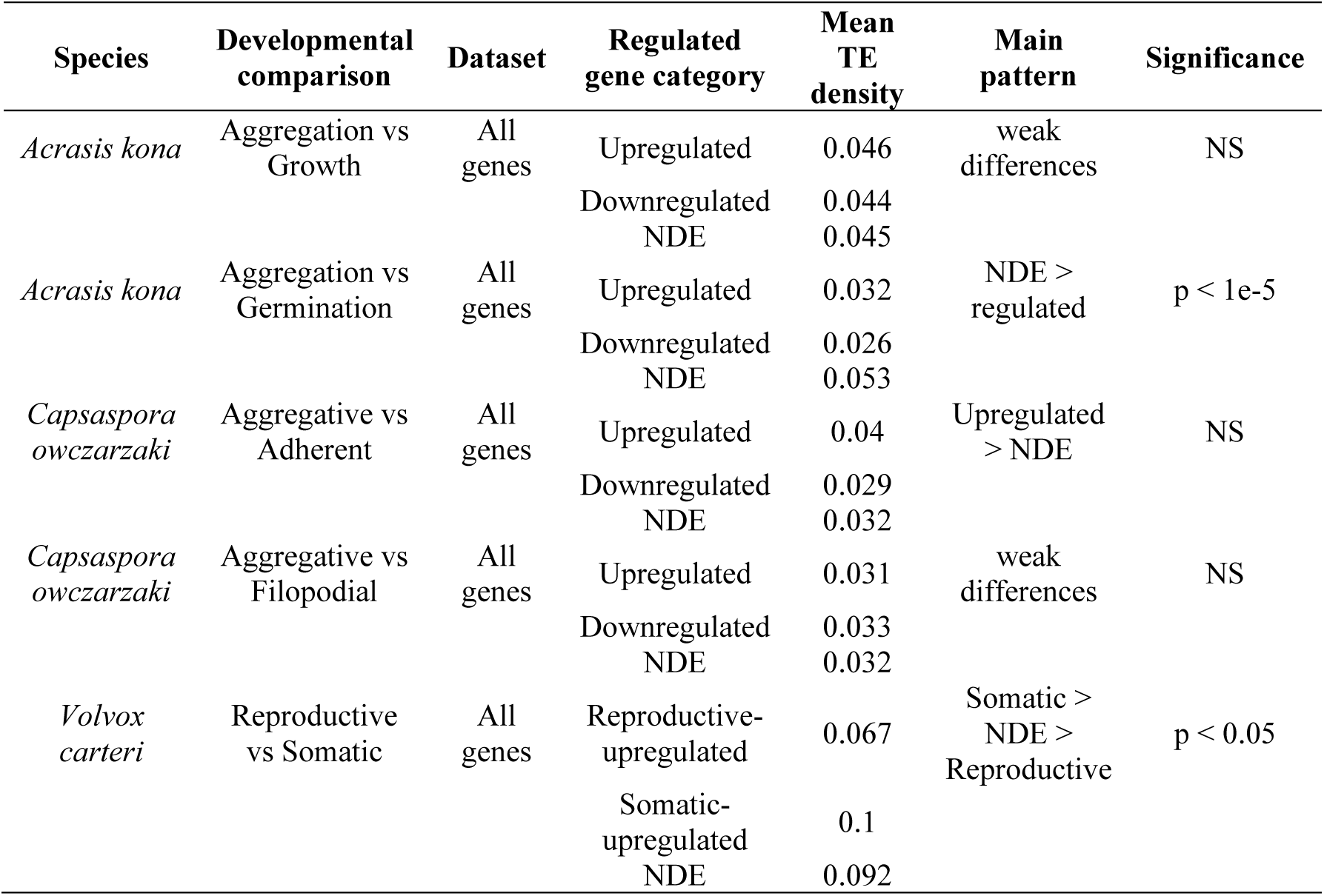
Upstream TE-density patterns across developmental regulatory comparisons in protist systems. Mean upstream TE-density values calculated within 5 kb windows surrounding genes belonging to distinct developmental regulatory categories in *Acrasis kona*, *Capsaspora owczarzaki*, and *Volvox carteri*. All analyses were performed using the complete set of annotated genes (“All genes” dataset). Comparisons include aggregation-associated transitions in *A. kona* and *C. owczarzaki* as well as reproductive versus somatic differentiation in *V. carteri*. For each comparison, the table reports the number of genes assigned to each regulatory category, mean TE-density values, overall TE-density pattern, and statistical significance. The results reveal lineage-specific relationships between TE density and developmental gene regulation, including TE depletion near regulated genes in *A. kona*, relatively weak and variable patterns in *C. owczarzaki*, and increased TE density near somatic-associated genes in *V. carteri*.

In *Acrasis kona*, aggregation versus growth comparisons exhibited minimal differences in upstream TE density among regulatory categories. At the 5 kb window, upregulated (0.046), downregulated (0.044), and non-differentially expressed (NDE) genes (0.045) displayed nearly identical mean TE-density values, and statistical analyses did not detect significant differences among groups. Similar patterns were observed across the 1 kb and 2 kb upstream windows, where TE-density values also remained highly comparable among regulatory categories and lacked significant statistical support (Supplementary Table S3).

In comparison, aggregation versus germination comparisons revealed a pronounced depletion of TE density surrounding genes important during developmental transitions. Upregulated and downregulated genes exhibited substantially lower TE-density values (0.032 and 0.026, respectively) than NDE loci (0.053), and these differences were consistently supported by parametric, nonparametric, and permutation-based statistical analyses (p < 1 × 10⁻⁵). Similar depletion patterns were observed across all evaluated upstream windows (1, 2, and 5 kb), although the strongest and most consistent statistical support was detected at the 5 kb window (Supplementary Table S3).

In *Capsaspora owczarzaki*, comparisons involving the aggregative state produced comparatively weak and inconsistent TE-density patterns. During aggregative versus adherent transitions, upregulated genes exhibited moderately elevated TE density (0.040) relative to NDE genes (0.032), whereas downregulated genes displayed lower values (0.029). Aggregative versus filopodial comparisons showed only minor differences among regulatory categories, with mean TE-density values remaining relatively similar across groups. None of the *Capsaspora* comparisons reached statistical significance, suggesting comparatively weak associations between TE density and developmental regulation in this lineage.

In *Volvox carteri*, reproductive versus somatic comparisons revealed a contrasting pattern relative to *A. kona*. Analyses including all genes showed that somatic-upregulated genes exhibited higher upstream TE density (0.100) than both reproductive-upregulated genes (0.067) and NDE loci (0.092) at the 5 kb window. These differences were supported by multiple statistical tests, including Mann–Whitney U, Welch’s t-tests, and permutation analyses (p < 0.05). However, analyses restricted to TE-positive genes (Supplementary Table S4) revealed substantially reduced effect sizes and weaker statistical support among regulatory categories, indicating that the observed signal is driven primarily by the presence or absence of nearby TEs rather than by large differences in TE density among already TE-associated loci.

Although several comparisons were statistically significant, the absolute differences in TE density among regulatory categories were relatively modest, generally differing by only a few percent of upstream sequence composition across the analyzed 5 kb windows. These results therefore suggest modest but consistent differences in genomic TE environments across developmental regulatory groups.

## Discussion

### TE proportion and diversity across protist genomes

Across the examined phylogenetic comparisons, species exhibiting aggregative, cooperative, or differentiated developmental states generally contained higher overall proportions of transposable elements than representative unicellular taxa. This pattern was especially pronounced in *Acrasis kona* and *Volvox carteri*, which exhibited substantially expanded TE landscapes relative to closely related heterolobosean and volvocine unicellular representatives. Statistical analyses across the examined lineage pairs further supported a significant overall enrichment of TEs among species exhibiting greater developmental complexity. These findings are broadly consistent with observations from plants and animals, where TE accumulation has been associated with increased genome plasticity, regulatory diversification, and long-term expansion of gene regulatory networks (Bourque et al., 2018; Elbarbary et al., 2016).

However, incorporation of the newly available holozoan genomes revealed a more nuanced evolutionary pattern than initially anticipated. Although *Capsaspora owczarzaki* exhibited elevated TE abundance relative to many protist genomes, both *Pigoraptor* species also contained comparatively TE-rich genomes despite lacking multicellularity. Importantly, *Pigoraptor* species have been reported to exhibit transient multicellular aggregations, coordinated feeding behaviors, and syncytium-like cellular stages associated with cooperative interactions among cells (Hehenberger et al., 2017). These observations suggest that some genomic features associated with increased TE accumulation may emerge prior to stable obligate multicellularity and may instead characterize lineages capable of facultative multicellular or cooperative behaviors, such as those observed in *Pigoraptor* and *Capsaspora*. Consequently, TE expansion may reflect broader increases in developmental, behavioral, or regulatory complexity across lineages representing intermediate stages of multicellular evolution. Further investigation of TE organization and regulatory architecture in *Pigoraptor* may therefore provide important insight into evolutionary transitions preceding more stable multicellular systems.

Despite clear differences in overall TE proportion, TE diversity analyses revealed no universal relationship between multicellularity and increased TE diversity. Instead, Shannon and Simpson indices demonstrated strongly lineage-specific patterns. Within Heterolobosea, *Acrasis kona* exhibited substantially greater TE diversity than the compared *Naegleria* species, whereas *Tetramitus jugosus* displayed relatively high diversity despite very low total TE abundance. In Chlorophyceae, the unicellular alga *Chlamydomonas reinhardtii* exhibited the highest overall TE diversity among all examined assemblies despite containing lower total TE abundance than *Volvox carteri*. Similarly, within Filasterea, TE diversity patterns reflected distinct lineage-specific TE histories rather than a simple relationship with developmental complexity. Together, these results indicate that TE expansion within protist genomes frequently reflects amplification of particular TE lineages rather than uniform diversification across all TE categories.

This interpretation is further supported by the TE compositional analyses. The PCA demonstrated that species frequently diverged through expansion of specific TE groups rather than through complete restructuring of their TE repertoires. For example, the large TE burden observed in *Volvox carteri* was driven primarily by LTR retrotransposons, whereas rolling-circle elements contributed strongly to the distinctive TE landscape of *Capsaspora owczarzaki*. Conversely, *Chlamydomonas reinhardtii* retained a comparatively balanced TE composition despite lower total TE abundance. These findings suggest that TE evolutionary trajectories differ substantially among protist lineages and are shaped by lineage-specific amplification, retention, and turnover dynamics rather than by a single conserved genomic response associated with multicellularity.

Interestingly, not all aggregative systems exhibited substantial TE expansion. Although *Fonticula alba* forms multicellular fruiting bodies, its genome remains comparatively TE-poor and compositionally similar to several unicellular taxa, although some underestimation of TE content may reflect assembly incompleteness or fragmentation. This result indicates that transitions toward cooperative or multicellular behaviors can occur through distinct genomic routes and do not require major TE proliferation. Together, these observations support a model in which TE accumulation frequently accompanies multicellular or aggregative lifestyles but is not itself a prerequisite for the evolution of multicellular organization.

### TE density and developmental regulation across protist systems

Comparative TE-density analyses revealed even stronger lineage-specific patterns. In *Acrasis kona*, upregulated and downregulated genes during aggregation versus germination transitions consistently exhibited relatively TE-depleted upstream environments compared to non-differentially expressed genes. Although the absolute differences in TE density were modest, these patterns were significant and remained consistent across multiple statistical approaches. Meanwhile, aggregation versus growth comparisons exhibited almost identical TE-density values among regulatory categories and produced no significant differences. This suggests that TE depletion is not uniformly associated with all developmental transitions in *A. kona*, but instead characterizes specific regulatory programs linked to aggregation and germination.

At first glance, the relatively low TE density surrounding regulated genes in *A. kona* may appear unexpected given extensive evidence showing that TEs can contribute to regulatory innovation in many eukaryotic systems (Chuong et al., 2017; Sundaram & Wysocka, 2020). However, even genes classified as TE-associated in *A. kona* remained only modestly enriched for nearby TE sequence in absolute terms, generally differing by only a few percent of upstream sequence composition. Thus, the observed patterns likely reflect subtle but evolutionarily meaningful differences in local genomic environments rather than extensive TE-driven restructuring of promoters or enhancers.

One possible interpretation is that genes requiring rapid and coordinated transcriptional responses during aggregation occupy comparatively TE-poor regulatory neighborhoods in order to preserve regulatory precision. Numerous studies across animals, fungi, and plants demonstrate that TE insertions near promoters or enhancers can alter transcription factor binding, chromatin accessibility, or local regulatory interactions (Feschotte, 2008; Bourque et al., 2018). Developmentally responsive genes may therefore experience stronger selective pressure inhibiting nearby TE accumulation because even modest regulatory perturbations could disrupt coordinated multicellular behavior. Within this framework, the TE-depleted regulatory landscapes observed in *A. kona* may reflect evolutionary constraints associated with maintaining robust aggregation-associated transcriptional programs. Future analyses examining gene family expansion and TE-associated duplication events may further clarify whether TEs contribute indirectly to multicellular evolution through the generation of novel gene copies and regulatory diversity.

Notably, *Capsaspora owczarzaki* exhibited comparatively weak and inconsistent TE-density organization across developmental states involving aggregation. Some comparisons showed slightly elevated TE density near upregulated genes, whereas others revealed only minimal differences among regulatory categories. None of these contrasts reached strong statistical significance. These results suggest that TE-associated regulatory architecture in *Capsaspora* may be substantially less structured than in *Acrasis*. These differences may reflect lineage-specific variation in regulatory organization and developmental gene expression programs across aggregative protists.

The most contrasting pattern was observed in *Volvox carteri*. Unlike *A. kona*, somatic-associated genes in *Volvox* exhibited comparatively elevated upstream TE density relative to reproductive-associated genes. Importantly, this signal was strongest when all genes were included and became substantially weaker when analyses were restricted to TE-positive loci. This indicates that the primary pattern in *Volvox* is driven largely by whether genes occur near TEs at all rather than by large differences in TE density once nearby TEs are already present. Although the observed differences remained modest in absolute magnitude, the consistent enrichment of TEs near somatic-associated genes suggests that differentiated multicellularity in volvocine algae may involve a distinct relationship between developmental regulation and genomic TE environments.

These results indicate that TE-associated regulatory organization is not evolutionarily conserved across protist developmental systems. Instead, different lineages appear to have evolved distinct genomic relationships with nearby TEs. In some systems, such as *Acrasis kona*, developmental regulation is associated with relative exclusion of TEs from upstream regulatory regions, whereas in others, such as *Volvox carteri*, developmental differentiation may tolerate or even coincide with comparatively TE-enriched genomic environments. Meanwhile, *Capsaspora owczarzaki* appears to occupy an intermediate condition characterized by comparatively weak TE-density organization.

Importantly, the TE-density differences observed across all systems remained relatively subtle in absolute magnitude, often differing by only approximately 2–4% of upstream sequence composition. Consequently, these results should not be interpreted as evidence of extensive promoter restructuring driven by TEs. Rather, they support a model in which relatively small shifts in genomic TE organization may contribute to broader differences in regulatory architecture when acting across thousands of genes over evolutionary timescales.

Collectively, these findings emphasize that the contribution of transposable elements to developmental evolution is highly context-dependent. In some lineages, TEs may primarily function as genomic elements that must be selectively excluded from sensitive regulatory loci, whereas in others they may become integrated into broader regulatory landscapes associated with cellular differentiation and developmental complexity. These contrasting patterns illustrate that the evolutionary relationship between TEs and multicellularity cannot be explained through a single universal model but instead reflects diverse lineage-specific trajectories of genome evolution and regulatory organization.

## Conclusion

This study combined comparative transposable element annotation with genome-wide analyses of TE density surrounding developmentally regulated genes across multiple protist lineages exhibiting distinct developmental and cellular organizations. Comparative analyses revealed that species exhibiting aggregative, cooperative, or differentiated developmental states generally contained higher overall TE proportions than representative unicellular taxa, although this trend was not universal across all lineages. TE diversity and TE composition additionally exhibited strong lineage-specific patterns, indicating that TE evolution in protists is shaped by distinct genomic trajectories rather than by a single conserved relationship with multicellularity.

Comparative TE-density analyses further demonstrated that relationships between TEs and developmental regulation differ substantially among protist systems. *Acrasis kona* exhibited reduced TE density near genes regulated during aggregation versus germination transitions, whereas *Volvox carteri* displayed comparatively elevated TE density near somatic-associated loci. At the same time, *Capsaspora owczarzaki* showed relatively weak and inconsistent TE-density organization across developmental states. Together, these findings indicate that TE-associated regulatory landscapes are highly lineage-specific and do not follow a universal pattern associated with multicellular evolution.

Overall, this work emphasizes the importance of integrating comparative genomics and transcriptomics to better understand how transposable elements contribute to genome organization, regulatory architecture, and the evolution of developmental complexity across diverse eukaryotic lineages.

## Supporting information

Fasta file for TE data base

Supplementary tables for TE families

## Acknowledgments

This work utilized computational resources provided by the Texas Tech University High Performance Computing Center (TTU HPCC). PromethION sequencing for *Acrasis kona* was supported by the National Science Foundation, Division of Environmental Biology, Biodiversity and Systematics Cluster (grant DEB-2100888). Additional institutional support was provided by the Department of Biological Sciences at Texas Tech University.

## Data Availability

The genome assemblies analyzed in this study were obtained from publicly available repositories, including NCBI GenBank, the European Nucleotide Archive (ENA), and associated BioProjects under the accession numbers reported in Table 1. The newly generated Acrasis kona genome assembly has been deposited in NCBI under BioProject accession PRJNA1447523, BioSample accession SAMN57028165, and Whole Genome Shotgun accession JBWXZU000000000.

Transcriptomic datasets used for differential gene expression analyses were obtained from publicly available repositories. Accession numbers for all RNA-seq datasets are provided in Supplementary Table S1. The curated transposable element classification dataset generated during this study is available as Supplementary Table S2, and the complete curated transposable element consensus library used for RepeatMasker analyses is available as Supplementary File S1.

## Conflict of Interest

The authors declare no competing interests.

## Notes

### Competing Interest Statement

The authors have declared no competing interest.

