## Supplementary tables for "Evaluating the potential role and contribution of transposable elements to the evolution of microbial multicellularity across the tree of eukaryotes"

**Supplementary Table S1. Transcriptomic datasets used for differential expression analyses.**

| **Species** | **Developmental condition / cell type** | **Database** | **Accession** | **Reference** |
| --- | --- | --- | --- | --- |
| *Acrasis kona* | Growth vs aggregation stages | SRA | SRR22861965 | Sheikh et al., 2024 |
| *Capsaspora owczarzaki* | Life cycle stages | SRA | PRJNA20341 | Sebé-Pedrós et al., 2013 |
| *Volvox carteri* | Somatic vs reproductive cells | ENA | ERP022669 | Matt & Umen, 2018 |

**Supplementary Table S3. TE-density comparisons across upstream windows in *Acrasis kona.***

Mean TE-density values and statistical comparisons across 1 kb, 2 kb, and 5 kb upstream windows for differentially expressed and non-differentially expressed genes in *A. kona*. Statistical support was evaluated using Mann–Whitney U tests, Welch’s t-tests, and permutation-based analyses.

| **Comparison** | **Window** | **Groups Compared** | **Mean Group 1** | **Mean Group 2** | **Mann–Whitney p** | **Welch t-test p** | **Permutation p** |
| --- | --- | --- | --- | --- | --- | --- | --- |
| aggregation_vs_growth | 1 kb | UP vs NDE | 0.0432 | 0.0434 | 0.9009 | 0.9543 | 0.9502 |
| aggregation_vs_growth | 1 kb | DOWN vs NDE | 0.0415 | 0.0434 | 0.2165 | 0.6481 | 0.6604 |
| aggregation_vs_growth | 2 kb | UP vs NDE | 0.0451 | 0.0439 | 0.7513 | 0.7237 | 0.7191 |
| aggregation_vs_growth | 2 kb | DOWN vs NDE | 0.0415 | 0.0439 | 0.1696 | 0.5383 | 0.5503 |
| aggregation_vs_growth | 5 kb | UP vs NDE | 0.0461 | 0.0448 | 0.2185 | 0.6528 | 0.652 |
| aggregation_vs_growth | 5 kb | DOWN vs NDE | 0.0437 | 0.0448 | 0.8891 | 0.7276 | 0.7258 |
| aggregation_vs_germination | 1 kb | UP vs NDE | 0.0295 | 0.0516 | 3.20E-14 | 6.88E-20 | <1E-5 |
| aggregation_vs_germination | 1 kb | DOWN vs NDE | 0.023 | 0.0516 | 2.87E-42 | 3.74E-38 | <1E-5 |
| aggregation_vs_germination | 2 kb | UP vs NDE | 0.0301 | 0.0522 | 5.46E-18 | 2.98E-23 | <1E-5 |
| aggregation_vs_germination | 2 kb | DOWN vs NDE | 0.0234 | 0.0522 | 1.56E-60 | 1.45E-45 | <1E-5 |
| aggregation_vs_germination | 5 kb | UP vs NDE | 0.0321 | 0.0525 | 3.08E-25 | 1.66E-27 | <1E-5 |
| aggregation_vs_germination | 5 kb | DOWN vs NDE | 0.0259 | 0.0525 | 3.51E-67 | 3.27E-55 | <1E-5 |

**Supplementary Table S4.** Mean upstream TE-density values calculated within 5 kb windows surrounding genes containing non-zero upstream TE density (“TE-positive genes only”) in *Acrasis kona*, *Capsaspora owczarzaki*, and *Volvox carteri*. The table summarizes the number of TE-positive genes assigned to each regulatory category, mean TE-density values, overall TE-density patterns, and statistical significance. Relative to analyses including all genes, TE-positive-only analyses generally produced weaker effect sizes and reduced statistical support, indicating that many observed genome-wide patterns are driven primarily by the presence or absence of nearby TEs rather than by large differences in TE density among already TE-associated loci.

| **Species** | **Developmental comparison** | **Window** | **Regulatory group** | **N genes** | **Mean TE density** | **Main pattern** | **Significance** |
| --- | --- | --- | --- | --- | --- | --- | --- |
| *Acrasis kona* | Aggregation vs Growth | 5 kb | Upregulated | 561 | 0.147 | Weak differences | NS |
|  |  |  | Downregulated | 419 | 0.148 |  |  |
|  |  |  | NDE | 6472 | 0.15 |  |  |
| *Acrasis kona* | Aggregation vs Germination | 5 kb | Upregulated | 962 | 0.126 | NDE > regulated | p < 1e-5 |
|  |  |  | Downregulated | 884 | 0.128 |  |  |
|  |  |  | NDE | 5606 | 0.158 |  |  |
| *Capsaspora owczarzaki* | Aggregative vs Adherent | 5 kb | Upregulated | 148 | 0.163 | Upregulated > NDE | NS |
|  |  |  | Downregulated | 361 | 0.106 |  |  |
|  |  |  | NDE | 1751 | 0.118 |  |  |
| *Capsaspora owczarzaki* | Aggregative vs Filopodial | 5 kb | Upregulated | 396 | 0.123 | Weak differences | NS |
|  |  |  | Downregulated | 431 | 0.124 |  |  |
|  |  |  | NDE | 1433 | 0.117 |  |  |
| *Volvox carteri* | Reproductive vs Somatic | 5 kb | Reproductive-upregulated | 2414 | 0.115 | Somatic > NDE > Reproductive | Reduced significance |
|  |  |  | Somatic-upregulated | 2475 | 0.156 |  |  |
|  |  |  | NDE | 3737 | 0.147 |  |  |
